# Peptide Antidotes to SARS-CoV-2 (COVID-19)

**DOI:** 10.1101/2020.08.06.238915

**Authors:** Andre Watson, Leonardo Ferreira, Peter Hwang, Jinbo Xu, Robert Stroud

**Affiliations:** Ligandal Inc., San Francisco, CA, USA; Diabetes Center, University of California San Francisco, San Francisco, CA, USA; Department of Surgery, University of California San Francisco, San Francisco, CA, USA; Department of Biochemistry and Biophysics, University of California San Francisco, San Francisco, CA, USA; Toyota Technological Institute, Chicago, IL, USA

## Abstract

The design of an immunogenic scaffold that serves a role in treating a pathogen, and can be rapidly and predictively modeled, has remained an elusive feat. **Here, we demonstrate that SARS-BLOCK™ synthetic peptide scaffolds act as antidotes to SARS-CoV-2 spike protein-mediated infection of human ACE2-expressing cells**. Critically, SARS-BLOCK™ peptides are able to potently and competitively inhibit SARS-CoV-2 S1 spike protein receptor binding domain (RBD) binding to ACE2, the main cellular entry pathway for SARS-CoV-2, while also binding to neutralizing antibodies against SARS-CoV-2. In order to create this potential therapeutic antidote-vaccine, we designed, simulated, synthesized, modeled epitopes, predicted peptide folding, and characterized behavior of a novel set of synthetic peptides. The biomimetic technology is modeled off the receptor binding motif of the SARS-CoV-2 coronavirus, and modified to provide enhanced stability and folding versus the truncated wildtype sequence. These novel peptides attain single-micromolar binding affinities for ACE2 and a neutralizing antibody against the SARS-CoV-2 receptor binding domain (RBD), and demonstrate significant reduction of infection in nanomolar doses. We also demonstrate that soluble ACE2 abrogates binding of RBD to neutralizing antibodies, which we posit is an essential immune-evasive mechanism of the virus. SARS-BLOCK™ is designed to “uncloak” the viral ACE2 coating mechanism, while also binding to neutralizing antibodies with the intention of stimulating a specific neutralizing antibody response. Our peptide scaffolds demonstrate promise for future studies evaluating specificity and sensitivity of immune responses to our antidote-vaccine. In summary, SARS-BLOCK™ peptides are a promising COVID-19 antidote designed to combine the benefits of a therapeutic and vaccine, effectively creating a new generation of prophylactic and reactive antiviral therapeutics whereby immune responses can be enhanced rather than blunted.

## MAIN

We developed and characterized four novel synthetic peptide inhibitors, SARS-BLOCK™, to serve as combined prophylactic, therapeutic, and immune stimulants against the novel coronavirus. In this study, we utilized computational techniques to design peptides that mimic the SARS-CoV-2 spike protein’s receptor binding domain (RBD) to ACE2 (Figures 1 & 2), while displaying neutralizing antibody binding motifs. These peptides were designed and simulated using SWISS-MODEL, PDBePISA, RaptorX, and Ligandal’s proprietary nanomaterials and design approaches. After synthesizing the peptides using solid-phase peptide synthesis (SPSS), we characterized their binding to ACE2 and neutralizing antibodies via biolayer interferometry (Figures 3 & 4, Table 1). Next, we infected cells expressing ACE2 with pseudotyped lentiviruses displaying the SARS-CoV-2 spike protein and expressing a luciferase reporter, and measured infection inhibition and toxicity following exposure to 7 concentrations (30nM - 20μM) of peptides (Figures 5 & 7). We compared these results to infection inhibition and toxicity mediated by 6 concentrations of soluble ACE2 (4nM - 1μM), 7 concentrations of SARS-CoV-2 receptor binding domain (RBD, 1nM - 1μM), and 5 concentrations of a potent neutralizing antibody against the SARS-CoV-2 RBD (6nM - 500nM) (Figures 6 & 7). At the highest concentrations measured (20μM), we observed that peptides 1, 4, and 6 completely inhibited infection without toxicity, while Peptides 5 and 6 exhibited up to ∼95% infection inhibition at 6.66μM and Peptide 5 exhibited statistically significant inhibition of infection with concentrations as low as 30nM (Figures 5–7). These data correspond to the biolayer interferometry results showing potent inhibition of viral RBD binding to ACE2 following exposure to SARS-BLOCK™ peptides.

**Figure 1.**
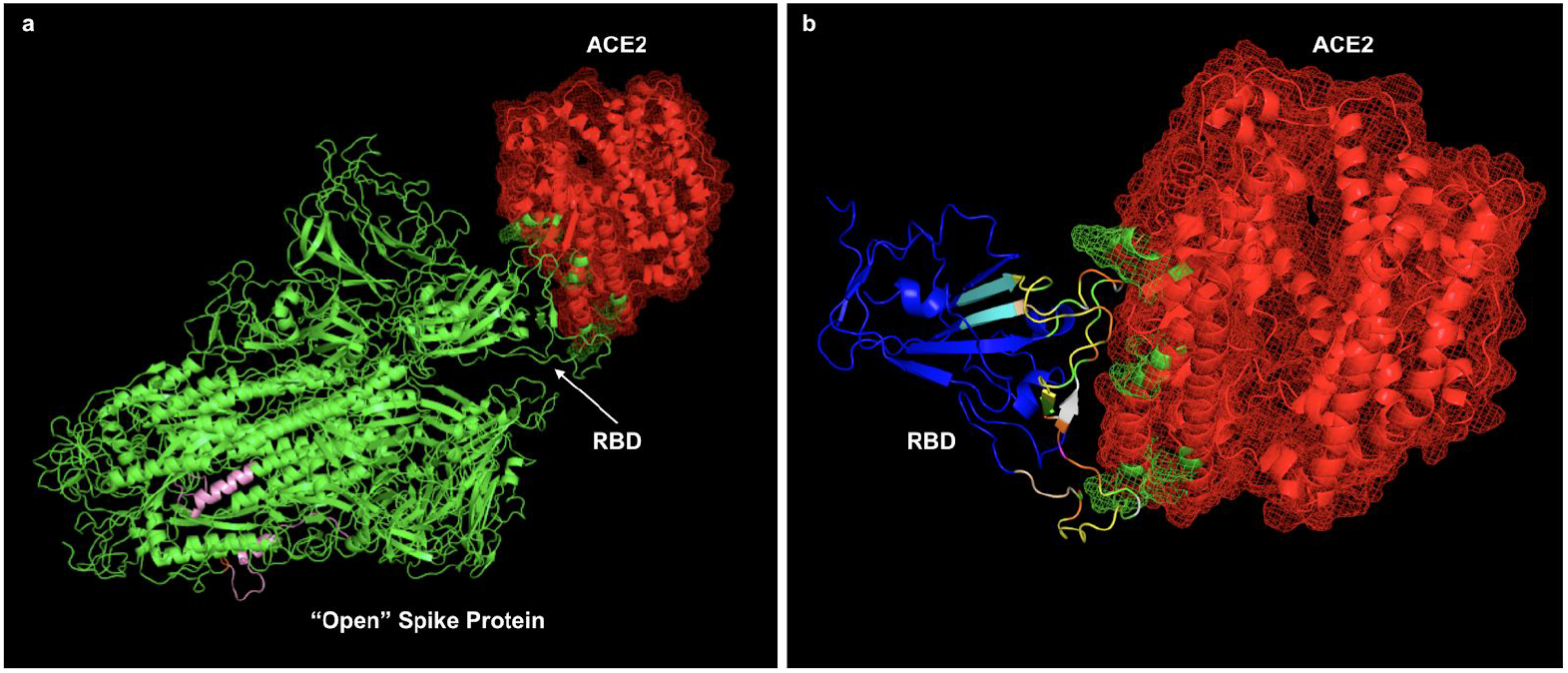
Initially, an “align” command was utilized in PyMOL with SARS-CoV-1 bound to ACE2 (PDB ID 6CS2) in order to approximate the binding interface of the SWISS-MODEL simulated SARS-CoV-2 (a). Selected MHC-I and MHC-II epitope regions for inclusion in Peptide 5 are colored pink and represent P807-K835 and A1020-Y1047 in the S1 spike protein, and were further refined by IEDB immune epitope analysis (a). Next, the receptor-binding domain of the SARS-CoV-2 S1 spike protein was truncated from the larger structure in its bound state to ACE2 (b). The resulting RBD structure was run through PDBePISA to determine interacting residues, highlighted in green on ACE2 (a and b). In b, green residues on the RBD indicate predicted thermodynamically favorable interactions between ACE2 and the S1 spike protein RBD, yellow indicate predicted thermodynamically neutral and orange indicate predicted thermodynamically unfavorable interactions. Cyan residues indicate the outer bounds of amino acids used to generate SARS-BLOCK™ peptides (V433 - V511). While the predicted binding residues did not overlap completely with subsequently empirically-validated sequences, the stretches of amino acids reflected in our motifs accurately reflected binding behavior (whereby N439, **Y449**, Y453, Q474, G485, **N487**, Y495, **Q498**, P499, and Q506 were suggested to be critical ACE2-interfacing residues by our PDBePISA simulation, while other groups’ subsequent mutagenesis studies determined that G446, **Y449**, Y453, L455, F456, Y473, A475, G476, E484, F486, **N487**, Y489, F490, Q493, G496, **Q498**, T500, N501, G502, and Y505 are critical for binding within the stretch of S425 - Y508).^35^ In sum, these residue predictions can be assessed as being precise, and accurate to within a few amino acids of actual binding behaviors—and represent a rapid and computationally minimalistic way to predict binding protein stretches without a structure when sufficiently long amino acid sequences are employed. Critically, we did not have to refine the amino acid sequences of SARS-BLOCK™ peptides to arrive at stretches of amino acids that were not empirically validated at the time of design.

**Figure 2.**
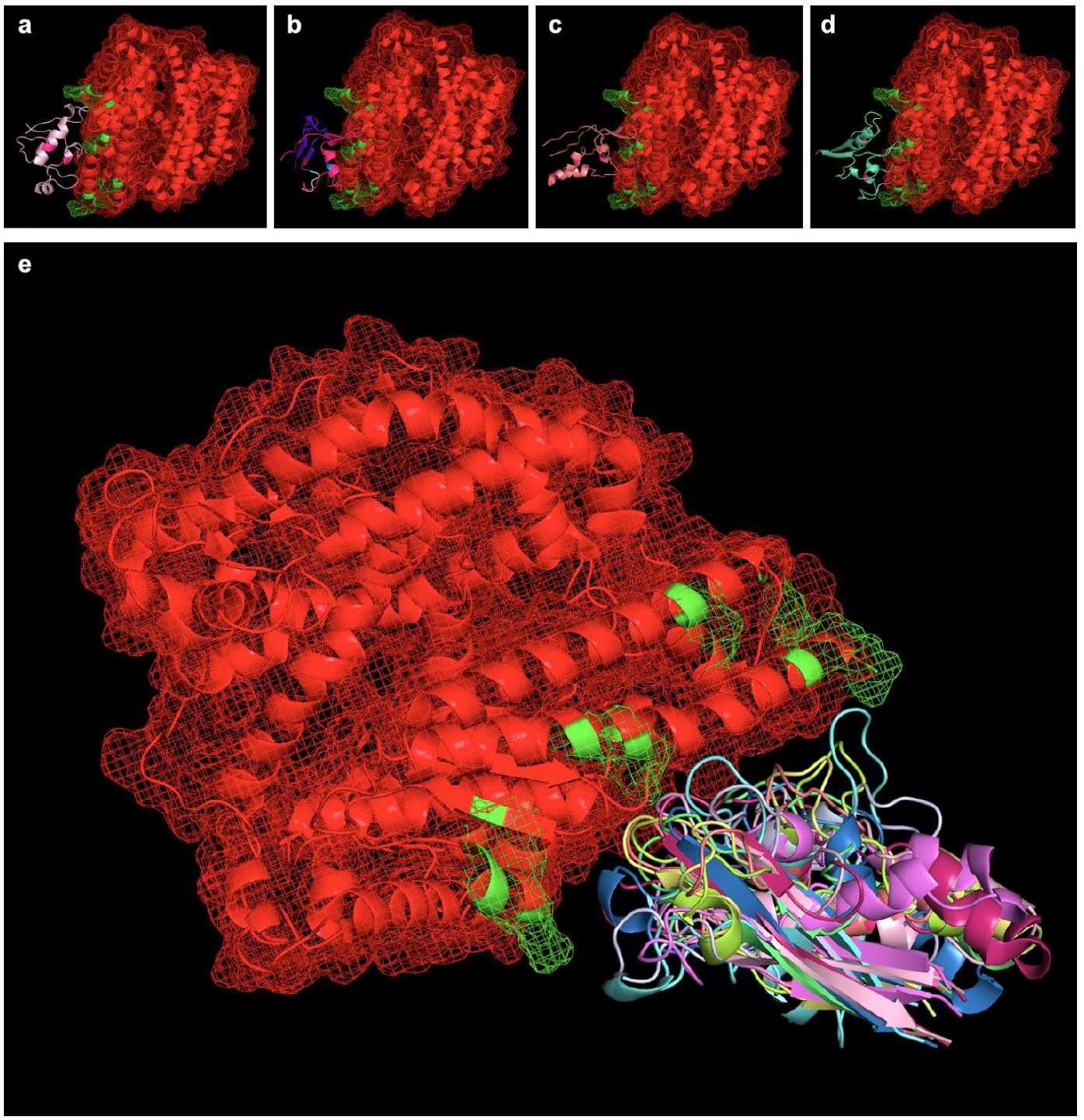
Peptides simulated via RaptorX were aligned with the SARS-CoV-2 binding interface of ACE2 (ACE2 in red, with PDBePISA-predicted binding interfaces in green). Shown from left to right (top) are SARS-BLOCK™ Peptides 1 (a), 4 (b), 5 (c), and 6 (d) bound to ACE2. Of note, all peptides exhibited two mutations introducing a disulfide bond to recreate the beta sheet structure of the SARS-CoV-2 receptor binding motif (RBM). Otherwise, Peptides 1 and 4 utilized the wildtype sequence, while Peptide 5 utilized MHC-I and MHC-II epitopes, and Peptide 6 utilized a GSGSG linker (white) in one of its non-ACE2-interfacing loop regions. Peptides 4, 5 and 6 exhibited additional, proprietary modifications to their sequences to facilitate appropriate folding, while Peptide 1 lacked this modification. Taking into account the 9 possible folded states generated for each peptide, we utilized PyMOL align commands which take into account multiple potential conformations of each peptide and may serve as a basis for future studies exploring more advanced molecular dynamics approaches for relaxing and simulating intramolecular interactions at the binding interface (e). In essence, the overlay of many possible folded states represents an electron distribution cloud of possible states that can be simulated for the minimal interfacial free energy of binding, and this approach requires vastly fewer computational resources than typically required for modeling binding pockets of *de novo* peptides or protein-protein interfaces without existing interfacial structures.

**Table 1.** Biolayer interferometry was used to determine dissociation constants (Kd) and RMax values (steady state binding analyses) for 1) SARS-BLOCK™ peptides 1, 4, 5 and 6 binding to ACE2, ACE2 and neutralizing antibodies binding to RBD, 3) peptides binding to neutralizing antibodies, and 4) RBD binding to antibodies in the presence of increasing concentrations of ACE2. Peptides 1, 4, 5 and 6 have dissociation constants of 1.8 ± 1.1μM, 5.2 ± 1.7μM, 2.4 ± 1.7μM, and 2.4 ± 1.7μM with ACE2, and 4.3 ± 1.0μM, 5.2 ± 1.7μM, unknown, and 3.3 ± 1.19μM with a neutralizing antibody, respectively. We were unable to determine Peptide 5 binding to the neutralizing antibody due to nonspecific interactions with the sensor tip. The dissociation constant of ACE2 with RBD is 2.3 ± 0.3nM, while the dissociation constant of the neutralizing antibody with RBD is 8.6 ± 1.1nM.

|  |  |  |  |
| --- | --- | --- | --- |
| RMax | $0.12111318 \pm 0.0239083$ | RMax | $0.22716674 \pm 0.0339753$ |
| KD | $1.80\text{E-}06 \pm 1.1\text{E-}06\text{M}$ | KD | $5.20\text{E-}06 \pm 1.7\text{E-}06\text{M}$ |
| **Peptide 1 Binding to RBD** |  | **Peptide 4 Binding to RBD** |  |
| RMax | $0.57363623 \pm 0.1333544$ | RMax | $0.13006174 \pm 0.032393$ |
| KD | $4.00\text{E-}06 \pm 2.2\text{E-}06\text{M}$ | KD | $2.40\text{E-}06 \pm 1.7\text{E-}06\text{M}$ |
| **Peptide 5 Binding to RBD** |  | **Peptide 6 Binding to RBD** |  |
| RMax | $0.71962807 \pm 0.0759471$ | RMax | $0.22716674 \pm 0.0339753$ |
| KD | $4.30\text{E-}06 \pm 1.0\text{E-}06\text{M}$ | KD | $5.20\text{E-}06 \pm 1.7\text{E-}06\text{M}$ |
| **Peptide 1 Binding to Neutralizing Antibody** |  | **Peptide 4 Binding to Neutralizing Antibody** |  |
| RMax | ∞ | RMax | $1.18656192 \pm 0.0552815$ |
| KD | $?$ | KD | $3.30\text{E-}06 \pm 3.9\text{E-}07\text{M}$ |
| **Peptide 5 Binding to Neutralizing Antibody** |  | **Peptide 6 Binding to Neutralizing Antibody** |  |
| RMax | $0.57847430 \pm 0.0155693$ | RMax | $4.40220793 \pm 0.159029$ |
| KD | $2.30\text{E-}09 \pm 3.0\text{E-}10\text{M}$ | KD | $8.60\text{E-}09 \pm 1.1\text{E-}09\text{M}$ |
| **ACE2 Binding to RBD** |  | **Neutralizing Antibody Binding to RBD** |  |

SARS-CoV-2, which causes COVID-19, is a global pandemic. At the time of this writing, more than 158,000 people have died from COVID-19 in the United States alone, and over 692,000 worldwide — corresponding to over 4.8M US and 18.2M global confirmed cases. The long-term health consequences of SARS-CoV-2 infection in recovered individuals remain to be seen, however include a range of sequelae from neurological to hematological, vascular, immunological, inflammatory, renal, respiratory, and potentially even autoimmune.^1,2,3,4,5,6,7,8,9,10^

These long-term effects are particularly concerning when factoring in the known neuropsychiatric effects of SARS-CoV-1, whereby 27.1% of 233 SARS survivors exhibited symptoms meeting diagnostic criteria for chronic fatigue syndrome 4 years after recovery. Furthermore, 40.3% reported chronic fatigue problems and 40% exhibited psychiatric illness.^11,12^ An affordable, globally deployable, room temperature stable, and repeatedly administrable therapeutic would address many of the risks of complications across the general population, and is urgently needed.

While numerous vaccines are in development, including Oxford’s and Moderna’s, key issues remain in assessing the long-term immune responses to both infections and vaccination. Additionally, the rapid spread and questionable long-term immunity towards the virus suggest that “herd immunity” strategies will both compromise population health and result in cumulative burden with associated long-term complications. At this time, consistent and persistent immune response across the entire population is speculative, and vaccine approaches have a likelihood of proving to be inadequate long-term solutions without immunomodulators and therapeutics to offset infection. Additionally, considerations of appropriate epitope display, glycosylations, T cell responses, antibody responses, innate immune responses, and immune evasive techniques of the virus must be taken into account across various demographics, both with and without vaccination.^13,14,15,16,17^ An ideal therapeutic strategy must take into account the root cause of infection, rather than attempting to treat specific downstream symptoms, given the measurable effects of various portions of the virus on extensive and specific cascades of physiological, intracellular and biological systems, as has been suggested with whole-interactome mass cytometry studies.^18^ SARS-CoV-2 has evolved numerous mechanisms for cloaking itself from the innate and adaptive immune systems.

To create a novel “antidote-vaccine” with potential utility as a prophylactic, immune-stimulant and therapeutic against the virus, we utilized several approaches to generate peptides modeled off the SARS-CoV-2 spike protein receptor-binding motif (RBM). We then performed binding and competitive association assays, as well as SARS-CoV-2 pseudotyped lentiviral infections of ACE2-expressing cells. We demonstrate that SARS-BLOCK™ peptides potently reverse ACE2 binding to viral spike proteins, with as much as ∼95-100% reduction in infection in cells expressing ACE2 receptors without causing toxicity (Figures 5–7). SARS-BLOCK™ prevents RBD binding to ACE2 (Figure 3) and binds to neutralizing antibodies with micromolar affinity (Figure 4a - 4d). We discover that soluble ACE2 potently inhibits binding of neutralizing antibodies (Figure 4g) to SARS-CoV-2 receptor binding domains, and that SARS-BLOCK™ peptides are also able to reverse this immune-evasive technique.

**Figure 3.**
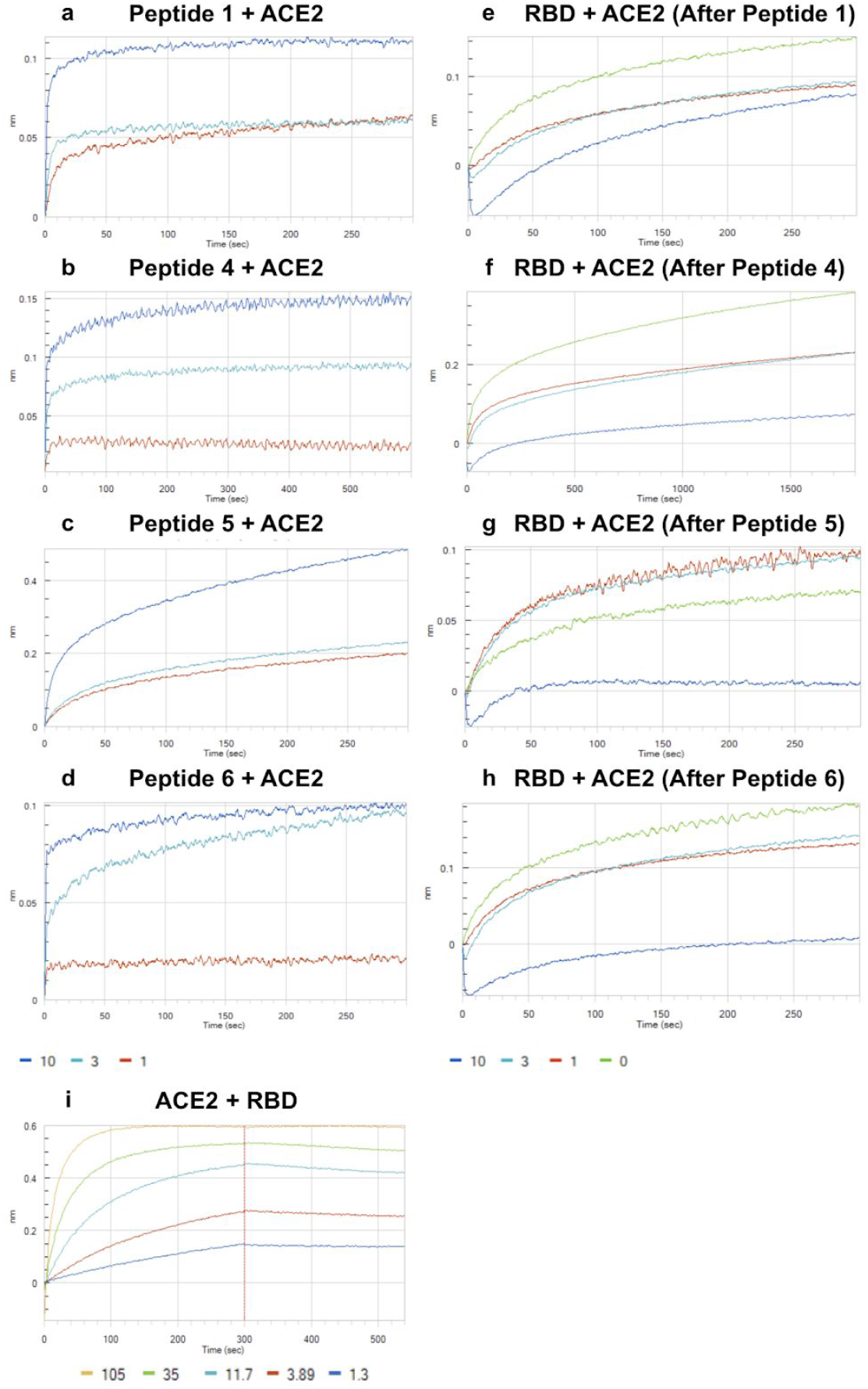
Biolayer interferometry was used to determine dissociation constants of SARS-BLOCK™ peptides associated with dimeric ACE2, and the inhibitory effects of peptides on ACE2 binding to RBD. All peptides exhibited potent inhibition of RBD binding to ACE2 at 10μM concentrations. Peptides were associated with ACE2 at 1, 3 and 10μM concentrations until saturation was observed (a-d). After peptide binding to ACE2, we measured ACE2 association of SARS-CoV-2 RBD at 35μM in the absence of peptides (e-h). Curiously, association of ACE2 with Peptide 5 at 1uM and 3uM enhanced RBD binding, while 10uM concentrations strongly abrogated binding (g). All other peptides exhibited a dose-response-like behavior in preventing RBD binding, including at 1 and 3μM concentrations (e,g,h). Of note, peptides were not included within the final solution of 35μM RBD, meaning that this assay was more of a measure of competitive irreversible antagonism versus IC50 determination. Finally, we compared these results to RBD-biotin captured on streptavidin sensor tips, and subsequently bound to monomeric ACE2 (i).

**Figure 4.**
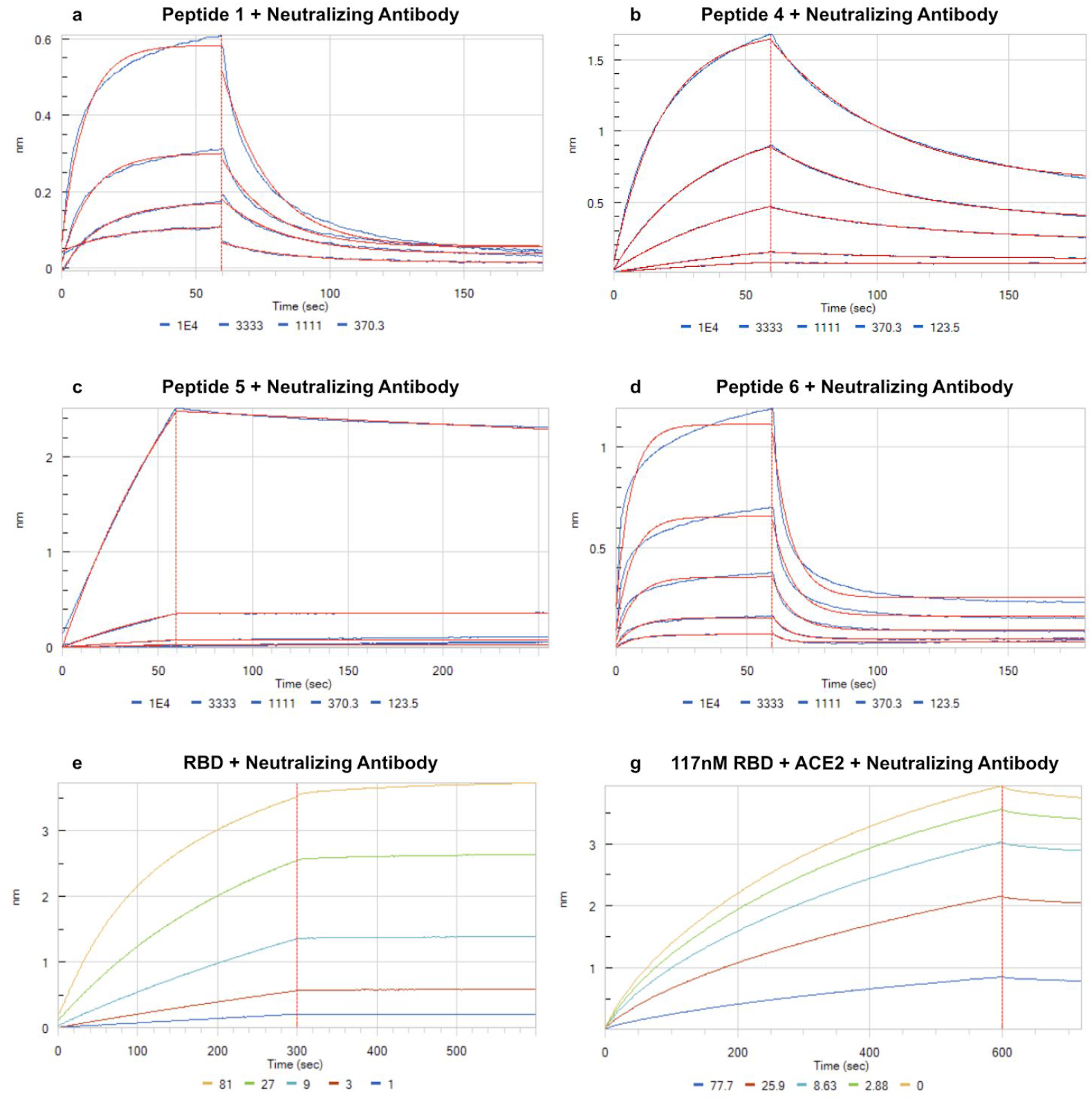
Biolayer interferometry was used to determine dissociation constants of SARS-BLOCK™ peptides associated with an IgG neutralizing antibody. Of note, Peptide 5 exhibited nonspecific binding to the sensor tip (c), preventing determination of Kd against the neutralizing antibody, and this was observed in all studies that did not utilize biotinylated substrates with biotin blocking of the sensor surface. However, we determined single-micromolar binding affinities for all other peptides with the neutralizing antibody (a,b,d). Next, we measured the dissociation constant of increasing concentrations of RBD with anti-RBD neutralizing antibody (e). Finally, 117nM RBD was mixed with increasing concentrations of ACE2 prior to introduction to immobilized neutralizing antibody, in order to demonstrate ACE2’s inhibition of neutralizing antibody binding to the RBD (g). The IC50 of ACE2 inhibiting interaction of RBD with the neutralizing antibody is interpolated to be ∼30-35nM for ACE2 when the RBD concentration is 117nM, confirming that ACE2 binds more potently to the RBD than the neutralizing antibody does, and that soluble ACE2 can act as a potent “cloak” against neutralizing antibody recognition even at fractional molarities to SARS-CoV-2 spike RBDs.

**Figure 5.**
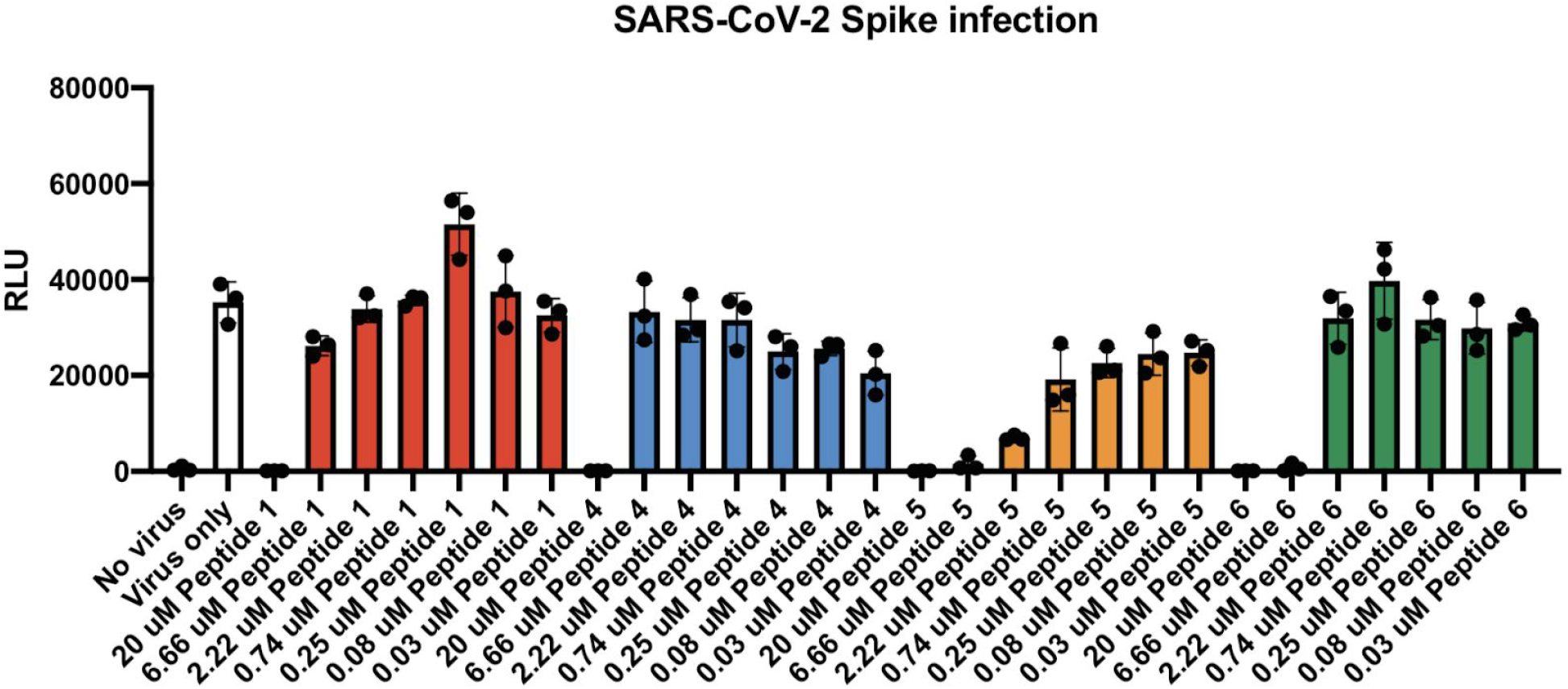
Peptides 1 and 4 did not block SARS-CoV-2 spike protein pseudotyped virus infection of ACE2-HEK293 cells at concentrations below 20μM, as assessed by luciferase activity 60h post-infection. Yet, both peptides 5 and 6 impeded viral infection at 6.66 micromolar, with Peptide 5 significantly exhibiting this blocking effect in the nanomolar range (80 nM and 30 nM, p < 0.05, t-test comparison with virus only). (*, p < 0.05; ***, p < 0.001; unpaired student’s t-test, technical triplicates). At 20μM, peptides 1, 4 and 6 achieved complete blockade of infection without statistically significant changes in toxicity vs. virus-only infected groups (luminescence values indistinguishable from uninfected controls).

In other words, we set out to develop a technology that could 1) block infection by inhibiting interaction of the SARS-CoV-2 spike protein with ACE2, as well as concomitant viral entry into ACE2-expressing cells, while 2) promoting binding of the virus and SARS-BLOCK™ peptides to neutralizing antibodies in order to stimulate a more efficacious immune response during infection. To this end, we also directly measured the binding of RBD and ACE2 with each other, as well as in the presence of SARS-BLOCK™ peptides (Figures 3 and 4), and discovered that soluble ACE2 forms an “immune cloak” upon the viral spike protein RBD, occluding the virus spike protein RBD from recognition by a potent (6nM Kd) neutralizing antibody against the RBD (Figure 4g). We also validated that SARS-CoV-2 RBD binds to ACE2 with 2.3±0.3 nanomolar affinity, in contrast to the whole spike protein in “open” conformation being reported to bind as strongly as ∼700 picomolar with ACE2, which we posit is a critical mechanism for immune evasion of the virus.^19^

ACE2, commonly known as the viral entry receptor for SARS-CoV-2, exists in both membrane-bound and soluble forms. In its soluble form, ACE2 potently prevents infection of ACE2-expressing cells *in vitro*, causing a ∼95% reduction in infection at ∼333.3nM and >98% reduction in infection at 1μM (Figure 6b). We observe statistically significant inhibition of SARS-CoV-2 pseudotyped lentiviral infections with ACE2 concentrations as low as 4nM, yet we also determine that soluble ACE2 prevents neutralizing antibodies from binding to the spike protein’s receptor binding domain (RBD). We posit that soluble ACE2 contributes to the immune cloaking and immune-evasive properties of the virus *in vivo*, essentially shielding the spike protein in its open conformation from recognition by the adaptive immune system.

**Figure 6.**
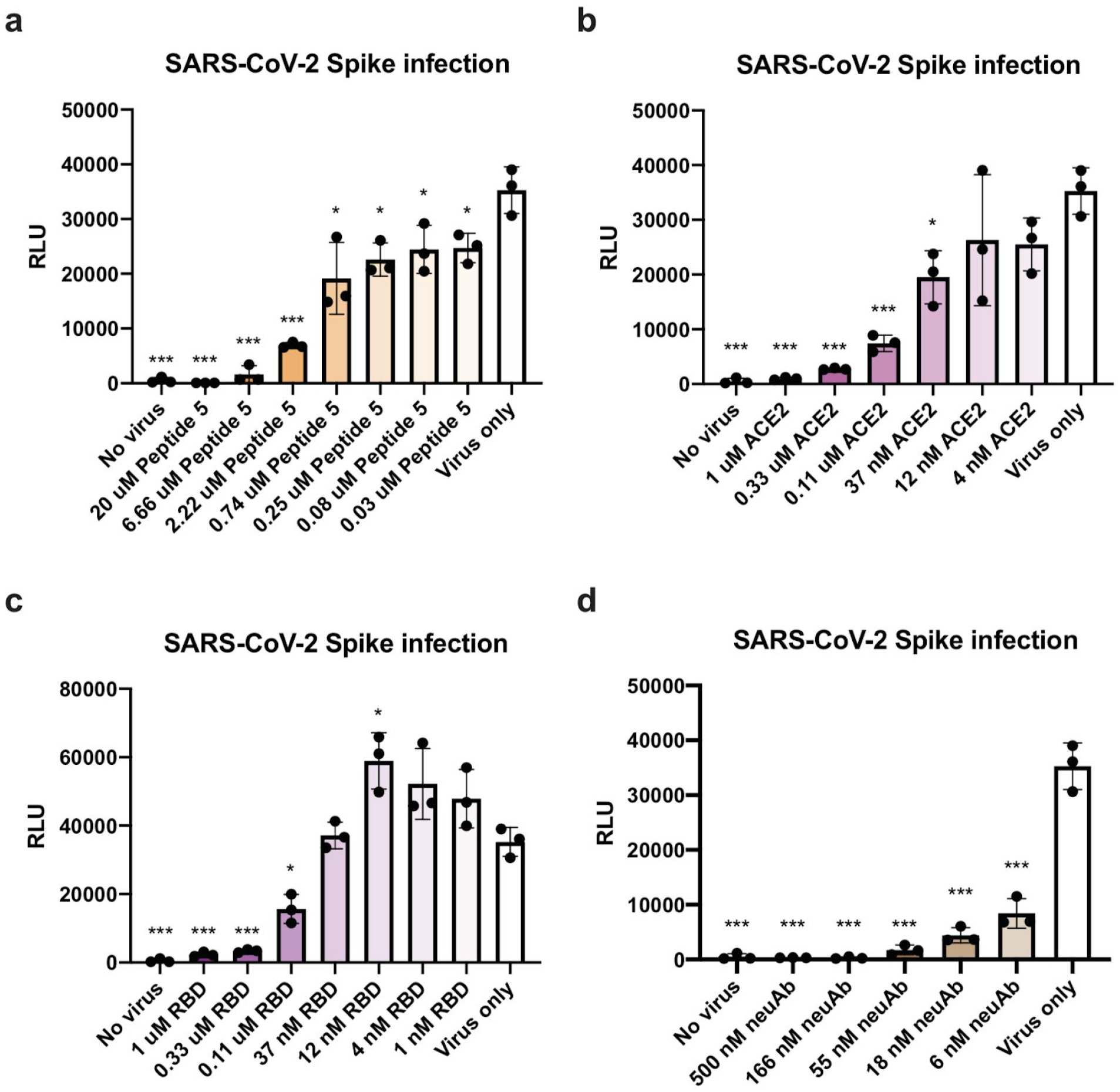
Peptide 5 exhibited an IC50 in the sub-micromolar range and an IC95 of ∼2.22uM. Of note, in parallel, we observed virtually complete inhibition of SARS-CoV-2 spike protein pseudotyped virus infection by soluble RBD and soluble ACE2 at 0.33uM, while a SARS-CoV-2 neutralizing antibody inhibited infection to a similar extent at concentration as low as 6nM. Intriguingly, 12nM RBD enhanced infection. (*, p < 0.05; ***, p < 0.001; unpaired student’s t-test, technical triplicates).

**Figure 7.**
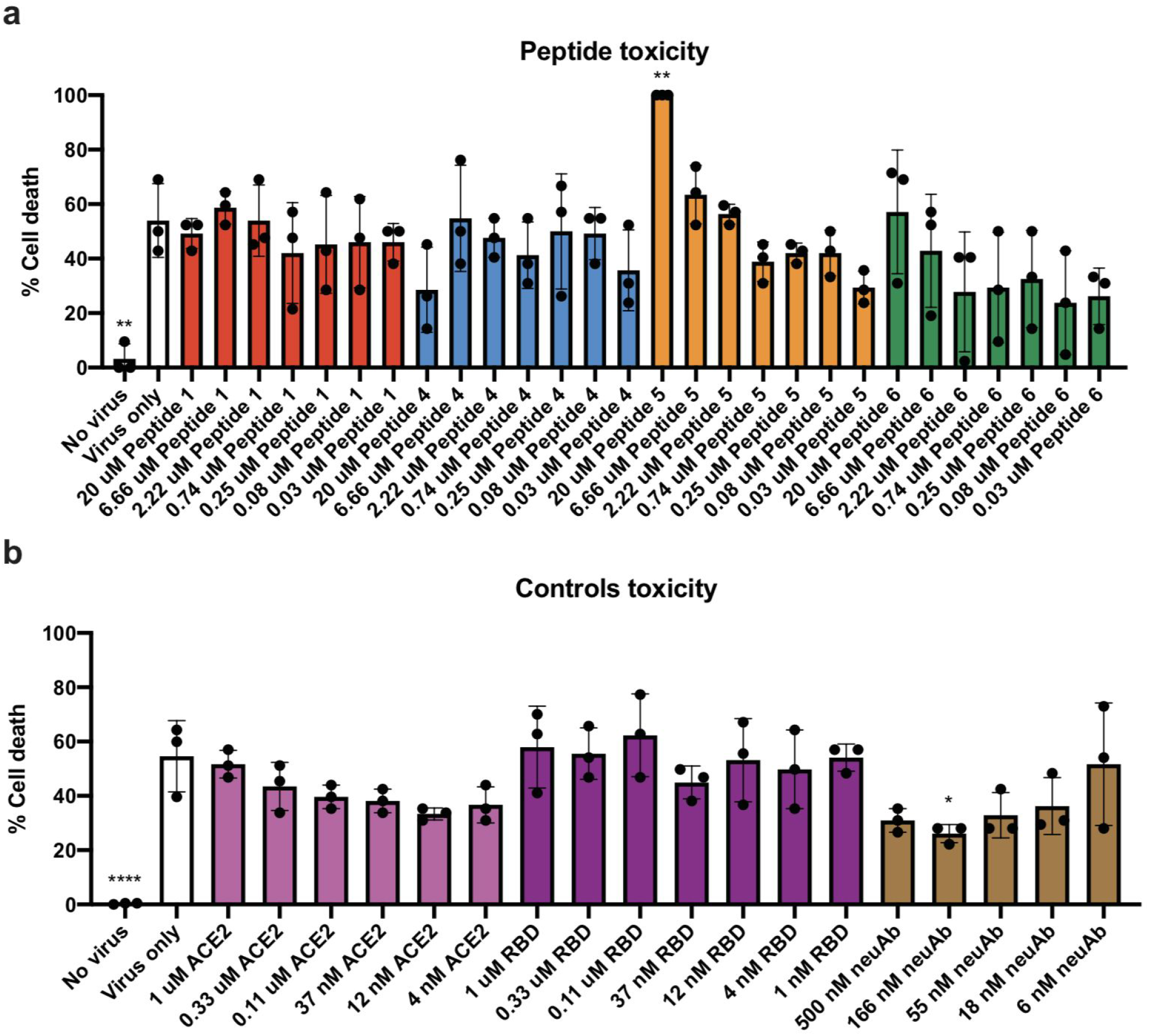
Importantly, with the exceptions of 20uM dose of Peptide 5 causing cell death (and leading to visible aggregation of the peptide in solution, due to poor aqueous solubility at this concentration), the addition of peptides, soluble ACE2, or soluble RBD at different concentrations did not result in statistically significant changes in cell viability in the presence of virus, ca. 50%. The SARS-CoV-2 neutralizing antibody exhibited a statistically significant (p < 0.05) survival-enhancing effect at 166nM.

In patients with heart failure, soluble ACE2 exists in plasma concentrations of 16.6 - 41.1ng/mL (1st and 4th quartile ranges), which corresponds to ∼193 - 478pM, while some studies report concentrations of 7.9ng/mL in acute heart failure patients and 4.8ng/mL in healthy volunteers, which corresponds to ∼92 and ∼56pM, respectively.^20,21^ Other studies report that male and female patients with type 1 diabetes (∼27.0 and ∼21.1ng/mL soluble ACE2, respectively) with comorbidities of diabetic nephropathy (∼27.7 - 30.2ng/mL and ∼19.9 - 21.5ng/mL, respectively), and/or coronary heart disease (∼35.5 and ∼27.0ng/mL, respectively) had higher circulating ACE2 concentrations than controls (∼25.6ng/mL and ∼20.3ng/mL, respectively), with higher arterial stiffness and microvascular or macrovascular disease also being positively associated with soluble ACE2 concentrations.^22^

In such ranges, it’s likely that ACE2 could enhance infection *in vivo* due to occluding the receptor binding domains of the S1 spike protein in open conformation, given that an individual virus spike only takes on this “open” conformation after exposure to furin (during biosynthesis) and TMPRSS2 (during membrane association).^23,24^ Additionally, the higher concentrations of ACE2 in patients with cardiovascular, diabetic, renal and vascular disease may further be associated with increased pathogenicity of SARS-CoV-2. Because of ACE2’s extremely potent binding affinity for SARS-CoV-2’s receptor binding domain, which we hypothesized would interfere with neutralizing antibody binding to the virus, the virus may avoid detection by the immune system as a function of soluble ACE2. SARS-CoV-2 viral titers in the blood of clinical specimens are lower relative to bronchoalvaeolar lavage, fibrobronchoscope brush biopsy, sputum, nasal swab, pharyngeal swab, and feces (an average of 2^4.6 reduction versus a cycle threshold of 30 corresponding to <2.6 × 10^4^ copies/mL), corresponding to ∼1000 viral copies per mL in the blood.^25^

Assuming ∼100 spikes per virus, this corresponds to ∼100,000 possible ACE2-binding sites per mL of blood if all spikes are in open conformation (∼1.66×10^−16^ M spike protein concentration in blood). However, given that the open conformation only occurs after TMPRSS2 cleavage, the starting position of each spike must be assumed as being closed, and likely only a fraction of these ∼100,000 sites are exposed for ACE2 or neutralizing antibody binding at any given point in time. Therefore, a ∼193 - 478pM soluble ACE2 concentration corresponds to 1.6×10^14^ −2.9×10^14^ molecules/mL (greater than one-billion-fold higher concentration than occupiable sites on the viral surface), which, when coupled to the ∼720pM - 1.2nM Kd of ACE2 to the spike protein in open conformation, suggests that SARS-CoV-2 would primarily exist with its “open” spikes occluded by ACE2 in blood. ACE2 is predicted to bind to certain SARS-CoV-2 RBD mutants with as little as 110 - 130pM Kd, and, importantly—when in fully “open” conformation—the SARS-CoV-2 spike proteins exhibit comparable binding affinity to neutralizing antibodies that compete for this same binding site.^19,26,27^ This is particularly troubling when considering the ability of ACE2 to hinder neutralizing antibody binding to this site, and that neutralizing antibodies are a product of B cell maturation, whereby B cells must mature antibodies and BCRs to reach single-digit nanomolar or picomolar binding affinities comparable in strength to ACE2-spike binding. It could be reasoned that any approach or endogenous mechanism that reduces neutralizing antibody binding to the SARS-CoV family spike proteins would result in less persistent B cell mediated immunity.

Indeed, SARS-CoV-2’s binding affinity to ACE2 is comparable to that of even potently neutralizing antibodies, and in our study we demonstrate that ACE2 severely abrogates antibody binding to the spike RBD as well as serving as potent inhibitor of infection of SARS-CoV-2 pseudotyped lentivirus in ACE2-expressing cells *in vitro*. Therefore, ACE2 serves both a protective function against infection and inhibitory function on immune recognition of the virus, acting as a competitive inhibitor of neutralizing antibody recognition against the spike protein (with binding affinities ranging from ∼676pM and 33.97nM).^28^

In our own studies, we found that the receptor binding domain (RBD) of SARS-CoV-2 spike bound to ACE2 with ∼2.3nM affinity, and that ACE2 could prevent association of a neutralizing antibody with the RBD that would otherwise have a ∼6nM binding affinity, even when ACE2 is presented in fractional concentrations to RBD (Figure 4). In sum, ACE2 binding to “open” conformation spike proteins is a viable mechanism at physiological ACE2 concentrations for inhibiting neutralizing antibody formation and binding against the spike protein RBD, and this bears further investigation as an *in vivo* neutralizing antibody avoidance technique of the virus.

The most recent spike protein mutation, D614G, seems to further increase the density of “open” spike proteins on the surface versus the original sequence, as well as the density of spikes in general, which notably makes this mutant likely to be more sensitive to neutralizing antibodies versus the aspartic acid (D) containing variant, while also increasing infectivity.^29,30^ In fact, the D614G variant seems to display ∼5x increased infectivity in ACE2-expressing cells with SARS-CoV-2 pseudotyped lentiviral infection assays.^31^ This likely relates to multivalent avidity increases associated with binding to cells expressing multiple membrane-bound ACE2 receptors, as is well characterized to be the case with viruses and other multivalent surface presentation substrates such as nanoparticles.^32,33,34^ As SARS-CoV-2 and COVID19 continue to ravage the world, it will be important to monitor the emergence and susceptibility of various mutants to “immune cloaking” by avoidance of neutralizing antibody recognition or recognition of the spike protein in “open” conformation, as well as any changes in avidity and affinity resulting from mutations at the binding interface or alterations in spike protein density.

Here, after modeling key interacting motifs of the spike protein and its receptor binding domain with ACE2 (Figure 1), we describe the simulation (Figure 2), design, synthesis and characterization (Figures 3 and 4, Table 1) of synthetic SARS-BLOCK™ peptides designed to block viral binding to cells expressing ACE2, prevent ACE2 association and subsequent cloaking of the viral spike protein, and stimulate an immune response due to binding to neutralizing antibodies and/or facilitating neutralizing antibody binding to the viral surface. Importantly, SARS-BLOCK™ is designed to expose the spike protein for recognition by the immune system, due to its potent inhibition of spike protein receptor binding domain (RBD) from binding to ACE2 (Figure 3). In summary, in contrast to neutralizing antibody therapies and other approaches that seek to target the virus, we developed a biomimetic virus decoy peptide technology that would compete for viral binding to cells, and expose the virus for binding to neutralizing antibodies.

## DISCUSSION

To our knowledge, this is the first instance whereby a synthetic peptide has been utilized to block a virus from associating with cells, while also including epitopes for antibody and T cell receptor formation. We demonstrate that SARS-BLOCK™ effectively blocks 95-100% of infection of pseudotyped lentiviruses displaying the SARS-CoV-2 spike protein infecting ACE2-expressing cells, and that SARS-BLOCK™ prevents association of the SARS-CoV-2 receptor binding domain (RBD) with ACE2 even at extremely high RBD concentrations (35μM).

Due to their binding to neutralizing antibodies against the RBD, SARS-BLOCK™ peptides are also expected to enhance immune response to SARS-CoV-2 rather than blunting it. In contrast, approaches such as ACE2-mimetic and antibody therapies are likely to reduce neutralizing antibody response to the virus, since they coat the virus and prevent binding of the adaptive immune system to the portion that is bound, which is the same segment of the spike protein necessary for B cell receptor (BCR) maturation into neutralizing antibodies targeting the spike protein RBD in its “open” conformation.

Importantly, SARS-BLOCK™ peptides are not expected to interfere in the activity of ACE2, due to binding to the face of the enzyme that does not metabolize angiotensin II. Critically for vaccine design and immune response promotion, these peptides are also designed to have modular epitopes for MHC-I and MHC-II recognition, which can be customized to the haplotypes of various patient populations, in addition to the inclusion of antibody-binding epitopes within the peptide sequences. Promisingly, recovered COVID-19 patients form dominant CD8+ T cell responses against a conserved set of epitopes, with 94% of 24 screened patients across 6 HLA types exhibiting T cell responses to 1 or more dominant epitopes, and 53% of patients exhibiting responses to all 3 dominant epitopes.^36,37^ Furthermore, patients with various HLA genotypes form MHC-I mediated responses to varying SARS-CoV-2 epitopes, and this can be predicted with bioinformatics approaches.^38^ While bioinformatic predictions of MHC loading corresponding to various HLA genotypes do not predictively reveal which peptide sequences will or will not be loaded, they do create a comprehensive overview of the possible state-spaces for empirical validation. SARS-BLOCK™ peptides are designed to display modular motifs for priming clonal expansion of selective TCR repertoires, which can be facilitated by sequencing of recovered patient TCR repertoires, insertion into these scaffolds, and finally assessing HLA genotypes of target populations, which we aim to assess in future studies.^39^ This affords a facilitated method for rapid vaccine and antidote design, coupling bioinformatics with structural and patient-derived omics data to create an iterative design approach to treating infectious agents.

Our future studies will also examine the role of altered epitope presentation and multivalent displays in creating enhanced antidote-vaccine technologies, while we demonstrate proof of principle for antibody recognition and effective viral blockade in this current study. Notably, using the approaches herein, we only synthesized four peptides, and all were “hit” compounds requiring minimal compute resources to design. The synthetic nature of SARS-BLOCK™ affords utility in tethering these peptides to a variety of multivalent substrates via click chemistry, which include but are not limited to C60 buckminsterfullerene, single and multi-walled carbon nanotubes, dendrimers, traditional vaccine substrates such as KLH, OVA and BSA, and the like — though we examine the bare alkyne-terminated peptides in this study. The synthetic nature, *in silico* screening, and precise conformation of these peptides allows for rapid synthesis without traditional limitations of recombinant, live-attenuated, gene delivery system, viral vector, and/or inactivated viral vaccine approaches. Due to the click chemistry nature of SARS-BLOCK™, it may also serve as a drug and gene delivery carrier by modifications with electrostatic sequences, or by click chemistry onto lipidic or other nanoparticles. We envision SARS-BLOCK™ and future permutations of these compounds vastly facilitating the design, development and scale-up of precise antidotes and vaccines against a variety of infectious agents as part of broader biodefense initiatives.

SARS-BLOCK™ peptides are designed to overcome many limitations associated with antibody therapies, ACE2-Fc therapies, and other antiviral therapeutics. Though neutralizing antibodies may be used as “stopgap” therapeutics to prevent the progression of disease, the transient nature of administered antibodies leaves the organism susceptible to reinfection. Furthermore, as we demonstrate in this study, ACE2 is a potent inhibitor of neutralizing antibody binding to the SARS-CoV-2 spike protein receptor binding domain, and this may relate to why short-lived antibody responses are a hallmark component of SARS-CoV-1 infections in many patients, whereby SARS-CoV-2 exhibits as much as 10-15x stronger binding to ACE2 than SARS-CoV-1 and would likely exhibit even stronger antibody maturation issues according to this hypothesis.^40,41,42,43^ Therapeutics that mimic ACE2 and shield this key epitope are likely to bias antibody formation towards off-target sites, which could contribute to antibody-dependent enhancement (ADE), vaccine-associated enhanced respiratory distress (VAERD), and a host of other immunological issues upon repeat viral challenge.^44,45,46,47,48,49,50,51,52,52^ With SARS-CoV-1, a marked lack of peripheral memory B cell responses was observed in patients 6 years following infection.^53^ These key issues are also important to consider in vaccine development, as there is precedent for enhanced respiratory disease in vaccinated animals with SARS-CoV-1.^54^ Thus, any approach that promotes a specific and neutralizing immune response, whether freestanding or in conjunction with another vaccine approach or infection, should be considered as an alternative to immunosuppressant and potentially off-target antibody forming approaches.

In particular, any approaches that have potential to limit endogenous antibody formation should be carefully reconsidered, due to the viral immune-evasive techniques already spanning a gamut of mechanisms, including but not limited to the S1 spike protein switching between “open” and “closed” conformations, heavy glycosylation limiting accessible regions, and also the presentation of T cell evasion due to MHC downregulation on infected cells and potential MHC-II binding of the SARS-COV-2 spike protein limiting CD4+ T cell responses, which all may be factors in contributing to T cell exhaustion and ineffective and/or transient antibody and memory B cell responses in infected patients.^55,56,57,58^ Indeed, severe and critically ill patients exhibit extreme B cell activation and, presumably, antibody responses.^59,60^ Yet, poor clinical outcomes are seen, suggesting that immune evasion and/or off-target antibody formation is dominant. The extent to which various factors individually play in contributing to these phenomena remains poorly understood. Surely, COVID-19 presents itself as a multifactorial disease with a cascade of deleterious effects. Also, the potential for reinfection across cohorts of varying disease severity remains to be fully elucidated, though numerous clinical and anecdotal reports indicate that immunity to coronaviruses is markedly short-lived, with seasonal variations in susceptibility to reinfection with alpha- and betacoronaviruses being frequently observed, and some antibody responses lasting for no longer than 3 months.^61^ With SARS-CoV-2 in particular, patients developing moderate antibody responses are seen to have undetectable antibodies in as little as 50 days.^62^ Additionally, one study on 149 recovered individuals reported that 33% of study participants did not generate detectable neutralizing antibodies 39 days following symptom onset, and that the majority of the cohort did not have high neutralizing antibody activity.^63^ In sum, any strategy that risks further contributing to these immune-evasive properties warrants caution. An ideal therapeutic strategy should enhance neutralizing antibody formation, not blunt it, while also preventing the virus from entering cells and replicating. In our study, we offer clues as to the role of soluble ACE2 in abrogation neutralizing antibody binding to the viral spike protein RBD, and a potential therapeutic solution to this immune evasion.

Importantly, we believe that SARS-BLOCK™ peptides will be useful as both antidotes and vaccines, due to the presence of key epitopes for antibody formation, and the performance of Peptide 5 (which exhibits one MHC-I and one MHC-II epitope) within these experiments suggests future utility for our multifunctional scaffolds in eliciting T cell responses against dominant viral epitopes, including ones that are not on the spike protein. The MHC-I and MHC-II domains can be flexibly substituted to match HLA types in various populations, or pooled across panels of peptides exhibiting multiple domains. Because SARS-BLOCK™ mimics the virus, rather than binding to it, and also due its its ability to displace ACE2 from cloaking the virus spike protein, we believe that these multifunctional peptides will prove to be an effective immune-enhancing strategy in infected patients, with additional potential to serve as a prophylactic vaccine.

Therefore, SARS-BLOCK™ peptides are an elegant solution towards preventing viral association with ACE2 and infection, while also contributing to a decrease in soluble ACE2 shielding of the virus. The thermodynamically favorable interaction of an antibody with the virus (∼6nM kD with the neutralizing antibody we studied) versus our peptides (∼1uM kD) suggests that the peptides can dissociate ACE2, promote antibody formation against the virus during infection, and preferentially train the immune system to eliminate the virus. Additionally, SARS-BLOCK™ effectively blocks infection by viruses displaying SARS-CoV-2 spike proteins. Our future studies will examine the improvements and therapeutic/vaccine utility of these peptides when displayed multivalently versus as free peptides, as well as refinements to structure and binding to further improve binding affinity and physiological behavior.

## MATERIALS AND METHODS

### SIMULATION AND DOCKING OF SARS-COV-2 SPIKE PROTEIN IN THE ABSENCE OF STRUCTURAL DATA

In order to elucidate the binding motif of the receptor binding domain, in the absence of structural data, it was necessary to utilize the results of prior crystallography experiments on SARS-CoV-1 with ACE2. SWISS-MODEL was utilized to generate a SARS-CoV-2 spike protein structure prior to the availability of Cryo-EM or X-ray crystallography data in February of 2020.^64,65,66,67,68,69^

The SARS-CoV-2 spike protein structure was aligned with SARS-CoV-1 spike protein bound to ACE2 (PDB ID 6CS2) using PyMOL (The PyMOL Molecular Graphics System, Version 2.3.5 Schrödinger, LLC.). This structure was then run through PDBePISA to determine the Gibbs free energy (ΔG) and predicted amino acid interactions between the SARS-CoV-2 spike protein and the ACE2 receptor.^70^ Upon the availability of structural data, this approach was compared and determined to have correctly identified the stretches of amino acids necessary for binding to ACE2.

### DESIGN AND SIMULATION OF SYNTHETIC PEPTIDES MIMICKING SPIKE PROTEIN RECEPTOR BINDING MOTIF

Following the simulation of structure and binding of the SARS-CoV-2 spike protein receptor binding domain (RBD), a truncated receptor binding motif (RBM) was gathered. This sequence was designed to recreate the structure of the larger protein in this motif, with key modifications performed to facilitate beta sheet formation. These modified peptide sequences were then simulated using RaptorX.

RaptorX is an efficient and accurate protein structure prediction software package, building upon a powerful deep learning technique.^71,72^ Given a sequence, RaptorX runs a homology search tool HHblits to find its sequence homologs and build a multiple sequence alignment (MSA), and then derives sequence profile and inter-residue coevolution information.^73^ Afterwards, RaptorX feeds sequence profile and coevolution information to a very deep convolutional residual neural network (of ∼100 convolution layers) to predict inter-atom distance (i.e., Ca-Ca, Cb-Cb and N-O distance) and inter-residue orientation distribution of the protein under prediction. To predict inter-atom distance distribution, RaptorX discretizes the Euclidean distance between two atoms into 47 intervals: 0-2, 2-2.4, 2.4-2.8, 2.8-3.2,…, 19.6-20, and > 20A. To predict inter-residue orientation distribution, RaptorX discretizes the orientation angles defined in into bins of 10 degrees.^74^

Finally, RaptorX derives distance and orientation potential from the predicted distribution and builds 3D models of the protein by minimizing the potential. Experimental validation indicates that such a deep learning technique is able to predict correct folds for many more proteins than ever before and outperforms comparative modeling unless proteins under prediction have very close homologs in PDB (Protein Data Bank).

For our studies, the peptides in their 9 possible folded states were overlaid with the SARS-CoV-2 receptor binding domain docked to ACE2 using PyMOL align commands to approximate binding, and then exported to PDBePISA to estimate binding pockets before synthesis and analytical characterization.

### IMMUNE EPITOPE MAPPING AND INCLUSION WITHIN PEPTIDES

IEDB is a predictive epitope discovery tool for determining possible MHC-I, MHC-II, and non-classical MHC restricted binding epitopes across various HLA genotypes.^75^ We utilized IEDB to predict key epitopes prior to clinical data emerging on various T cell receptor (TCR) responses across populations with various HLA alleles, and compared epitopes on SARS-CoV-1 with known immunogenicity to predicted and conserved epitopes for MHC-I and MHC-II response on SARS-CoV-2.^76^ A key region of SARS-CoV-1 (LPDPLKPTKRSFIEDLLFNKVTLADAGFMKQYG) was defined based on known immunogenicity of the monovalent peptide in terms of its ability to elicit an MHC-I response and antibody response, whereas many other peptides were only immunogenic while present multivalently. Next, a known MHC-II domain from SARS-CoV-1 was also defined (ASANLAATKMSECVLGQSKRVDFCGKGYH). These two peptides were compared to the stretches in SARS-CoV-2, and shown to have a high degree of homology in SARS-CoV-2, with sequences QILPDPSKPSKRSFIEDLLFNKVTLADAGFIK (804-835) and ASANLAATKMSECVLGQSKRVDFCGKGY (1020-1047). IEDB determined that sequences KMSECVLGQSKRV and LLFNKVLTA of SARS-CoV-2, representing MHC-II and MHC-I binding domains for HLA-A*02:01, respectively, would be immunogenic with percentile ranks of 0.9 and 1.2, respectively. These sequences were included within a non-interfacing loop structure of Peptide 5, whereas Peptide 4 included a GSGSG linker and Peptide 6 included the wildtype receptor binding motif (RBM) sequence for the loop. Other epitopes may be flexibility included within these modular stretches of Peptide 5 and homologues, representing 9- and 13-mer amino acid sequences corresponding to MHC-I and MHC-II binding, respectively.^77,78^

### PEPTIDE SYNTHESIS

The best performing *in silico* peptides were synthesized via microwave-assisted solid-phase peptide synthesis (SPSS) by a commercial manufacturer, sb-PEPTIDE (France). Mass spectrometry was utilized to confirm the appropriate peptide molecular weights.

### BIOLAYER INTERFEROMETRY

Biolayer interferometry is a label-free method for measuring the wavelength shift of incident white light following loading of a ligand upon a sensor tip surface, and/or binding of soluble analytes to that ligand or the sensor tip surface.^79^ The wavelength shift corresponds to the amount of analyte present, and can be used to determine dissociation constants and competition between multiple analytes and the immobilized ligand. An Octet® RED384 biolayer interferometer (Fortebio) was utilized with sensor tips displaying anti-human IgG Fc (AHC), streptavidin (SA), nickel-charged tris-nitriloacetic acid (NTA), or anti-penta-his (HIS1K) in 96-well plates. For streptavidin tips, we utilized 1mM biotin to block the surface after saturation with a given immobilized ligand. After protocol optimization with his-tagged vs. biotin-tagged variants of ACE2 and RBD, we determined that peptide analytes in solution exhibited nonspecific binding to the sensor tip surface with NTA and HIS1K tips, whereas biotinylated surfaces minimized this non-specific binding. Furthermore, ACE2-his (Sino Biological) and RBD-his (Sino Biological) exhibited extremely weak binding to HIS1K tips, so we opted to utilize dimeric-ACE2-biotin (UCSF) and RBD-biotin (UCSF) on SA tips, as well as neutralizing monoclonal IgG antibody against the SARS-CoV-2 spike glycoprotein (CR3022, antibodies-online) on AHC tips for all studies. Non-specific binding was still observed with Peptide 5 binding to a neutralizing antibody on AHC tips, which complicated efforts of determining the Kd of the Peptide 5 analyte vs. the neutralizing antibody ligand. All stock solutions were prepared in 1X PBS containing 0.2% BSA and 0.02% Tween20. Next, the following ligands and analytes were studied:

1. Dimeric ACE2-biotin was immobilized on SA tips (∼2.5nm capture).

a. Peptides 1, 4, 5 and 6 were introduced to immobilized ACE2 in concentrations of 1, 3 and 10μM (Figure 3a - 3d).
b. Sensor tips were removed from peptide solutions and introduced to 35μM RBD-his (Sino Biological) (Figure 3e - 3h).
2. RBD-biotin was immobilized on SA tips (∼5nm capture).

a. ACE2-his (Sino Biological) was introduced to immobilized RBD in concentrations of 1.3, 3.9, 11.7, 35 and 105μM (Figure 3i).
3. Neutralizing IgG antibody was immobilized on AHC tips (∼1nm capture).

a. Peptides 1, 4, 5 and 6 were introduced to immobilized ACE2 in concentrations of 0.37, 1.11, 3.33 and 10μM (Figure 4a - 4d).
b. RBD-his (Sino Biological) was introduced to immobilized neutralizing antibody (CR3022, antibodies-online) in concentrations of 1, 3, 9, 27 and 81μM.
c. 117nM RBD-his (Sino Biological) was mixed with ACE2-his in concentrations of 0 (RBD-only), 2.88, 8.63, 25.9, and 77.7μM, and then introduced to immobilized neutralizing antibody (CR3022, antibodies-online).

### INFECTION OF ACE2-HEK293 WITH SARS-COV-2 SPIKE PROTEIN PSEUDOTYPED LENTIVIRUS

ACE2-HEK293s (BPS Bioscience) were cultured and transduced in opaque 96-well white plates (Corning®) with pseudotyped lentivirions displaying the SARS-CoV-2 spike glycoprotein (BPS Bioscience). A neutralizing monoclonal IgG antibody against the SARS-CoV-2 spike glycoprotein (CR3022, antibodies-online), ACE2 (Sino Biological), receptor-binding domain (RBD) of spike glycoprotein (Sino Biological), and SARS-BLOCK™ peptides (Ligandal) were used as inhibitors of infection. Infection was quantitated via bioluminescence, and toxicity was characterized via a trypan blue absorbance assay utilizing a Synergy™ H1 BioTek spectrophotometer 60h following viral transduction. The cells were stained with trypan blue for 15 minutes, washed 3x with ice-cold calcium- and magnesium-containing 1x PBS, and lysed before transfer to a clear-bottom plate for absorbance measurements as described elsewhere.^80^

## AUTHOR CONTRIBUTIONS

AW prepared the manuscript, performed crude *in silico* docking of RaptorX-simulated peptides, and conceived and designed SARS-BLOCK™ peptides as well as the antibody and TCR epitope designs. LF performed cell culture and lentiviral transduction of HEK-ACE2 cells in the presence of peptides, antibodies, RBD, and ACE2. AW performed spectrophotometry and trypan blue assays on the transduced cells. LF and AW performed data processing and analysis of spectrophotometry results of bioluminescent and absorbance experiments. LF prepared figures on lentiviral transduction luminescence and performed statistical analysis. PH and AW performed biolayer interferometry assays. PH prepared figures on biolayer interferometry and performed statistical analysis. JX performed RaptorX modeling. RS reviewed the manuscript and provided feedback on the experimental procedures.

## ACKNOWLEDGEMENTS

We thank the Jim Wells lab (UCSF) for providing biotinylated dimeric ACE2 and RBD for biolayer interferometry studies. This research was supported by private funding by Ligandal Inc.

